# Diverse expression and modification of tRNAs and tRNA-derived RNAs in 19 mouse tissues

**DOI:** 10.1101/2025.11.11.685198

**Authors:** Aidan C. Manning, Patricia P. Chan, Todd M. Lowe

## Abstract

Transfer RNAs (tRNAs) and their derived small RNAs (tDRs) vary in abundance and processing across mammalian tissues. However, these differences and the regulatory mechanisms controlling them are poorly understood. Dysregulation of tRNAs has increasingly been implicated as a critical factor in disease progression; thus, a deeper understanding of the homeostatic variance across somatic tissues is vital. In this study, we applied Ordered Two-Template Relay sequencing (OTTR-seq) to examine small RNAs in 19 tissues from adult mice, quantifying mature tRNAs and tDRs at single-transcript resolution and inferring select base-modification signatures from characteristic reverse-transcriptase-mediated misincorporation. We observed tissue-biased expression of many unique transcripts and identified tDRs with greater tissue specificity than their parent tRNAs. Differences between tRNAs with the same anticodon but containing variations elsewhere in the sequence (isodecoders) suggest diverse regulatory processes across tissues that alter modification, translation functionality, stability, and tDR synthesis. These data provide an isodecoder-resolved reference for tissue-biased tRNA and tDR regulation in mice and form a framework for exploring disease-associated remodeling of the tRNA pool.

## Introduction

Transfer RNAs (tRNAs) play a central role in ribosomal translation. tRNA genes are also important chromosomal regulatory elements, acting as powerful insulators or enhancers or nearby genes, depending on their genomic context (Raab et al. 2012). Over the past decade, it has been demonstrated that their activity can vary by cell type (Isakova et al. 2020; Kapur et al. 2024) or in response to changing cellular conditions (Goodarzi et al. 2016; Liu et al. 2016; Aharon-Hefetz et al. 2020). tRNA-derived small RNAs (tDRs; also known as tRFs, tiRNAs, and tsRNAs) are generated from the cleavage or fragmentation of mature tRNAs, thereby expanding the regulatory repertoire of tRNAs beyond their primary role in translation (Shen et al. 2018; Boskovic et al. 2020a). Recent work has attributed these tDRs as critical factors in maintaining cellular homeostasis (Kim et al. 2019; Boskovic et al. 2020b), stress response (Fricker et al. 2019; Li et al. 2022), and disease progression pathways (Goodarzi et al. 2015; Shao et al. 2017; Vaidhyanathan et al. 2025). As tRNAs are increasingly recognized as directly modulating numerous cellular processes, a deeper understanding of their steady-state abundance and modification patterns across diverse cellular contexts is important.

The best-known example of tissue-specific tRNA expression and regulation is tRNA-Arg-TCT-4-1 in the brain, where the loss of this transcript is associated with altered translation initiation, activation of the integrated stress response, and suppression of mTOR signaling (Ishimura et al. 2014; Kapur et al. 2020). Only a small number of other tRNAs have been identified as having highly biased tissue expression using specialized tRNA-sequencing methods (Pinkard et al. 2020; Gustafsson et al. 2025). Comparatively, tDRs have been more widely characterized across different somatic tissues, mainly through secondary inclusion in miRNA sequencing libraries. For example, tDRs have been shown to be differentially expressed across tissues, such as the 5′ tDR of tRNA^Glu(UUC)^, which is enriched in the pancreas (Isakova et al. 2020). However, these analyses typically identified the source of tDRs at the level of tRNA isoacceptor (only anticodon is known) rather than mapping to nucleotide-resolution unique transcripts. None of these studies has information on tDR modifications. Due to these limitations, an integrated baseline view of both tRNA and tDRs across a broad range of somatic tissues is notably absent.

Studying tRNAs is technically challenging due to their highly stable structure and dense RNA modifications, averaging thirteen per molecule in mammals (Schimmel 2018). Multiple high-throughput sequencing methods, including ARM-seq, DM-tRNA-seq, mim-tRNA-seq, Nano-tRNA-seq, PANDORA-seq, and QuantM-tRNA-seq, have been developed to specifically focus on the quantification of tRNAs or tDRs, with some applications including RNA modification detection (Cozen et al. 2015; Zheng et al. 2015; Behrens et al. 2021; Lucas et al. 2024; Shi et al. 2021; Pinkard et al. 2020; Clark et al. 2016). However, none of these methods were designed to capture the expression of individual tRNA and tDR transcripts, with RNA modification information, making it difficult to assess and understand the relationship between these molecules within cells. Although liquid chromatography-tandem mass spectrometry (LC-MS/MS) is the gold standard for identifying RNA modifications, its technical limitations, cost, and requirement for relatively large input material amounts preclude its application at scale for most research groups. Alternate approaches include employing different chemical treatments such as sodium bisulfite (Xue et al. 2020), sodium borohydride (Lin et al. 2019), and CMCT (Marchand et al. 2020) in combination with high-throughput RNA sequencing to detect specific types of RNA modifications (m^5^C, m^7^G, and pseudouridine, respectively, for these examples). Yet these methods each yield information on only one type of modification, with most focused on sparsely modified mRNAs.

Here, we employed a robust sequencing method, OTTR-seq (Upton et al. 2021), that, unlike previous tRNA-focused approaches, captures the expression of densely modified tRNAs and tDRs without the need for enzymatic pre-treatments, and provides the identification of RNA modification sites through reverse transcription (RT) misincorporations at single-nucleotide resolution. In tandem with our tRNA-seq data analysis pipeline, we conducted a comprehensive assessment of the homeostatic variance of mature tRNAs and their corresponding tDRs, including RT-sensitive modification data across a panel of nineteen mouse tissues. We confirm prior observations of known tRNA-biased expression and explore a wealth of new evidence with immediate implications for human health and disease.

## Results

### Small RNAs exhibit distinct tissue-based expression signatures

To better understand the natural variance of tRNAs and derived small RNAs, we profiled their abundance in concert with other small ncRNAs across a panel of 19 mouse tissues from three adult female C57BL/6J mice using OTTR-seq (Upton et al. 2021) (Figure 1A). The distribution of sequencing reads revealed that tRNAs (including pre-tRNAs, mature tRNAs, tDRs, and mitochondrial tRNAs) were the largest class of detected transcripts across nearly all tissues, comprising approximately 50-70% of the total small RNA pool (Figure 1B, top panel). The remaining reads mapped to other RNA classes, including ribosomal RNA (rRNA) fragments, microRNAs (miRNAs), small nucleolar RNAs (snoRNAs), small nuclear RNAs (snRNAs), fragments from messenger RNA (mRNA) transcripts, and unannotated regions (intergenic). The number of distinct small RNA molecules detected varied considerably across tissues, with skin and mammary gland showing the highest diversity; in contrast, brain and kidney exhibited the fewest unique transcripts (Figure 1B, bottom panel). The length distribution of tRNA-mapping reads revealed two distinct size profiles: mature tRNAs (approximately 73-76 nucleotides [nt]) and shorter tRNA-derived RNAs (tDRs) in the 15-25/30-35 nt ranges. For well-studied miRNAs, reads characteristically peaked at ∼20 nt, corresponding to the mature transcripts rather than precursor hairpins (Figure 1C). SnoRNA and snRNA reads included both full-length mature transcripts and fragments of diverse sizes (Figure 1C), with highly heterogeneous tissue-specific length profiles.

**Figure 1.**
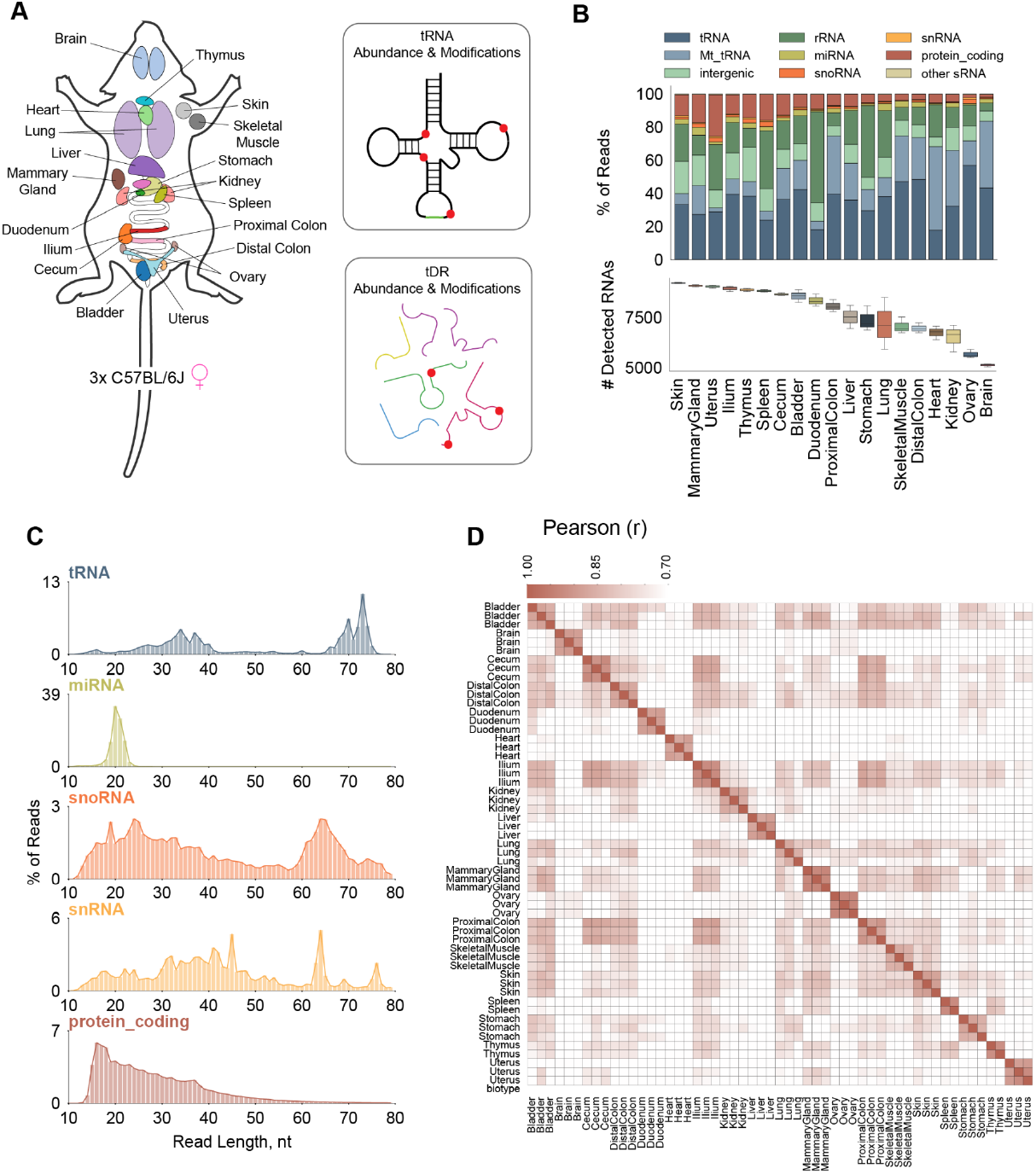
The small RNA landscape across mouse tissues. **(A)** RNA was isolated from 19 mouse tissues from three adult female C57BL/6J mice. Colors denote the different tissues assessed. **(B)** Composition and diversity of the small RNA pool across tissues. The top panel shows the relative abundance (% of reads) of different small RNA classes in each tissue. The bottom panel displays the number of unique small RNA transcripts detected in each tissue, sorted by median value. **(C)** Characteristic read length distributions for major small RNA classes. Histograms show the size distribution of reads mapping to tRNAs, miRNAs, snoRNAs, snRNAs, and protein-coding transcripts, aggregated across all samples. **(D)** Heatmap of pairwise Pearson correlation (r) for global small RNA expression profiles across all samples. The clustering reveals that samples from the same or functionally related tissues have more similar small RNA signatures.

An assessment of overall small RNA expression between tissues revealed that those with related physiological functions often shared similar small RNA profiles. For instance, distinct clusters were observed for digestive tract components (cecum with proximal and distal colon; duodenum with ileum) and lymphoid tissues such as the spleen and thymus. These data collectively highlight the tissue-specific nature of small RNA populations and reflect their unique metabolic requirements and regulatory processes.

### Tissue comparisons reveal distinctly expressed tRNA isodecoders

To assess tRNA diversity across tissues, we analyzed sequencing data for 214 unique tRNA transcripts from the high-confidence tRNA gene set in the Genomic tRNA Database (GtRNAdb) (Chan and Lowe 2016). Interestingly, only 147 (68.7%) unique tRNA transcripts were expressed in at least one adult tissue; however, the number varied tissue-by-tissue (median 136 unique tRNAs; Figure 2A). We hypothesize that the remaining undetected subset may be under more specialized regulatory control, restricted to developmental stages, cellular stresses, or tissues we did not measure. We paid particular attention to tRNAs with the same anticodon but slightly different sequences (isodecoders) to determine whether additional, more recently evolved tRNA “versions” exhibit more specialized regulation. Specifically, the brain harbored the most diverse tRNA pool, while the mammary gland had the least, highlighting tissue-specialized regulation of these molecules.

**Figure 2.**
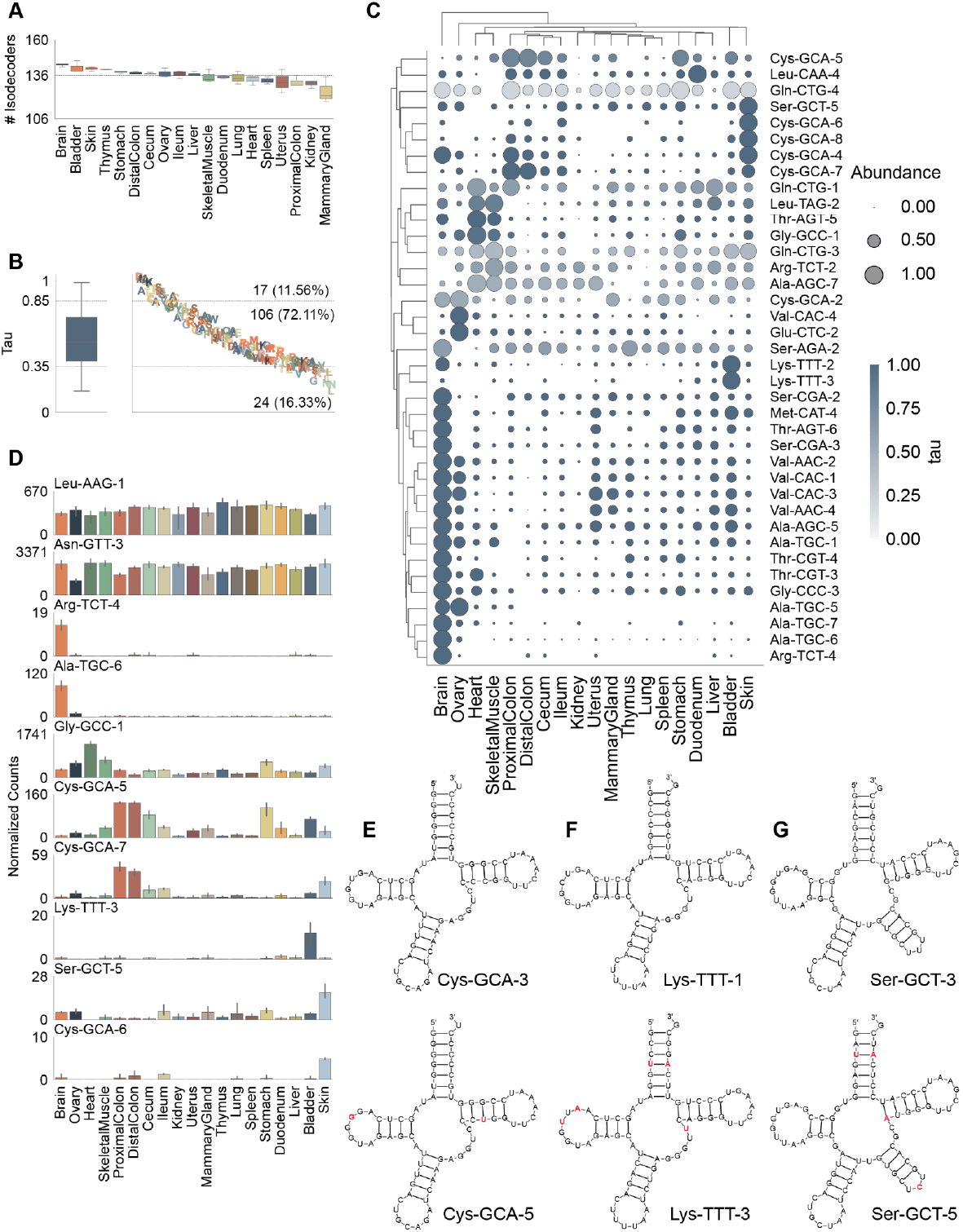
Isodecoder-level analyses of tissue-biased tRNAs. **(A)** The number of abundant tRNA isodecoders detected in each of the mouse tissues. **(B)** Classification of tRNAs based on tissue-specificity using the tau statistic. The box plot shows the distribution of tau scores, and the scatter plot categorizes individual tRNAs as tissue-unbiased (tau < 0.80), tissue-biased (0.8 ≤ tau < 0.95), or tissue-specific (tau ≥ 0.95). **(C)** Hierarchically clustered dot plot of the most significantly tissue-biased tRNAs. Dot size is proportional to the isodecoder’s abundance within a tissue, and color corresponds to its tissue-specificity score (tau). **(D)** Expression profiles for selected tissue-biased tRNA isodecoders. Bar plots show the normalized counts for each tRNA across the 20-tissue panel. **(E-G)** Comparison of secondary structures between highly abundant (‘major’) and tissue-biased (‘minor’) isodecoders. Nucleotide differences in the minor isodecoder compared to the major one are highlighted in red. **(E)** Colon-biased tRNA-Cys-GCA-5 compared to the major isodecoder tRNA-Cys-GCA-3. **(F)** Bladder-biased tRNA-Lys-TTT-3 compared to the major isodecoder tRNA-Lys-TTT-1. **(G)** Skin-biased tRNA-Ser-GCT-5 compared to the major isodecoder tRNA-Ser-GCT-3.

By comparing the expression values across the studied tissues, we identified 136 (92.52% of 147) differentially expressed tRNA isodecoders (adjusted p-value < 0.05), suggesting that nearly all expressed tRNAs were regulated at tissue level. We then applied tau statistics (Kryuchkova-Mostacci and Robinson-Rechavi 2017) and categorized these isodecoders as “tissue-specific” (17 tRNAs; 11.56%), “tissue-biased” (106 tRNAs; 72.11%), and “unbiased” (24 tRNAs; 16.33%) (Figure 2B). Clustering of the most significant tissue-biased tRNAs revealed distinct abundance patterns (Figure 2C), with the brain harboring the largest set of enriched isodecoders. The most tissue-specific tRNA in our samples is the well-studied brain-specific tRNA-Arg-TCT-4 (tau=0.99) (Figures 2C and 2D), one of the five tRNA^Arg(UCU)^ isodecoders with a mutation that increased seizure threshold and altered synaptic transmission when combined with impaired ribosome rescue factor GTPBP2 (Ishimura et al. 2014; Kapur et al. 2020). Consistent with the previous study using QuantM-tRNA-seq (Pinkard et al. 2020), we also found three tRNA^Ala(UGC)^ isodecoders (tRNA-Ala-TGC-5, 6, 7 with tau=0.95, 0.98, 0.97 respectively) and tRNA-Thr-AGT-6 (tau=0.90) among the top brain-specific tRNAs (Figure 2C). Moreover, our data reveal additional novel brain-biased tRNAs including tRNA-Ser-CGA-2 and tRNA-Ser-CGA-3 (Figure 2C). Yet, we noticed that some of the tRNAs differentially expressed in the central nervous system (CNS) tissues of the QuantM-tRNA-seq study were biased in other tissues. For example, tRNA-Ala-AGC-1 was thymus-biased while tRNA-Ser-GCT-5 was skin-biased (Figure 2G). This could be caused by the lack of the corresponding tissues in the previous study. Overall, 29 tRNAs (19.7%) had the highest abundance in the brain with the majority matching those enriched in the CNS tissues of QuantM-tRNA-seq study.

In addition, tRNAs tended to have similar expression biases in physiologically similar tissues, such as the skeletal muscle and heart, as well as different segments of the colon, including the distal, proximal, cecum, and the ileum. Among the colon-biased tRNAs were tRNA-Cys-GCA-5 and tRNA-Cys-GCA-7, which are both “minor” isodecoders, making up only ∼1.9% and 0.4% respectively of the total tRNA^Cys(GCA)^ pool. Unexpectedly, only two nucleotides in the sequence of tRNA-Cys-GCA-5 are different from the most abundant isodecoder tRNA-Cys-GCA-3. One of them is U16 at the D-loop of tRNA-Cys-GCA-3, a position that is possibly modified as dihydrouridine (Yu et al. 2011). Having a G instead indicates that tRNA-Cys-GCA-5 will likely follow an alternate regulation pattern due to the missing of dihydrouridine. The other change is from a C51:G63 pair in the T-stem to a U51:G63 pair, which causes a difference in the thermodynamics of the tRNA structure and is typically found in eukaryotic tRNA^Phe^ and tRNA^Tyr^ (Westhof et al. 2022) (Figure 2E). Other highly tissue-biased tRNAs included tRNA-Gly-GCC-1 in skeletal muscle and heart. Interestingly, the bladder harbored some of the most tissue-specific tRNAs, tRNA-Lys-TTT-2 and Lys-TTT-3, both of which were nearly absent across all other tissues (Figures 2C and 2D). The abundance of the bladder-biased tRNA-Lys-TTT-3 leads to approximately 15% shift of the tRNA^Lys(UUU)^ pool for this tissue. Instead of having the mammalian conserved C4:G69 pair (Westhof et al., 2022) as the most abundant isodecoder tRNA-Lys-TTT-1, tRNA-Lys-TTT-3 harbors a distinct U4:A69 in addition to sequence variations at positions 15 (G->A), 17 (C->U), and 48 (C->U) (Figure 2F). tRNA^Lys(UUU)^ typically has m^5^C48 modification, which serves as a critical regulator for transcript stability, translational efficiency, and tRNA processing (Blanco et al. 2014; Song et al. 2022). The change from C to U in tRNA-Lys-TTT-3 indicates the lack of this modification that suggests alternate regulation or functional mechanisms. Taken together, these tissue-biased shifts towards tRNA isodecoders with variability at known tRNA regulatory elements suggest differentiated mechanisms in tissues to modulate their tRNA pool.

### Modifications vary between tRNA isodecoders

Cytosolic tRNA modification profiles inferred by RT misincorporations in the sequencing alignments were generated as previously described (Zhang et al. 2025) to identify multiple modified bases, including m^1^A9, m^2^_2_ G26, ms^2^t^6^A37, and m^3^C47d, among others (Figure 3A). Using this methodology, we identified modified bases in 155 unique transcripts (Figure 3B), substantially expanding the set of known modified bases in mouse tRNA transcripts (Cappannini et al. 2024). Among these transcripts, at least one RNA modification was detected, with the majority having 3 to 4 detectable modifications (Figure 3C). Similar to human tRNAs, modifications in tRNA^Thr^ and tRNA^Tyr^ were more detectable (Zhang et al. 2025) across the mouse tissues while those in tRNA^Lys^ and tRNA^Gln^ were less detectable. For tRNAs with A34 at the anticodon, we found that all transcripts were edited to harbor an inosine (Figure 3B), consistent with the wobble pairing rule (A, C, U in mRNAs can be recognized and decoded by tRNAs with I34) (Agris 1991; Grosjean et al. 2010). Interestingly, almost all tRNAs having G at position 37 or C at position e2 (or 47d in the variable loop) were modified to m^1^G /wybutosine (yW) or m^3^C respectively (Figure 3B). Yet, some modifications were only detected in specific tRNAs. For example, only tRNA^Phe(GAA)^ harbored m^1^A at position 14 although no known enzyme is responsible for its deposition (Oerum et al. 2017). These data highlight that local nucleotide context, tRNA secondary and tertiary structure, or the expression of the modifying enzymes may play critical roles in regulating tRNA modifications.

**Figure 3.**
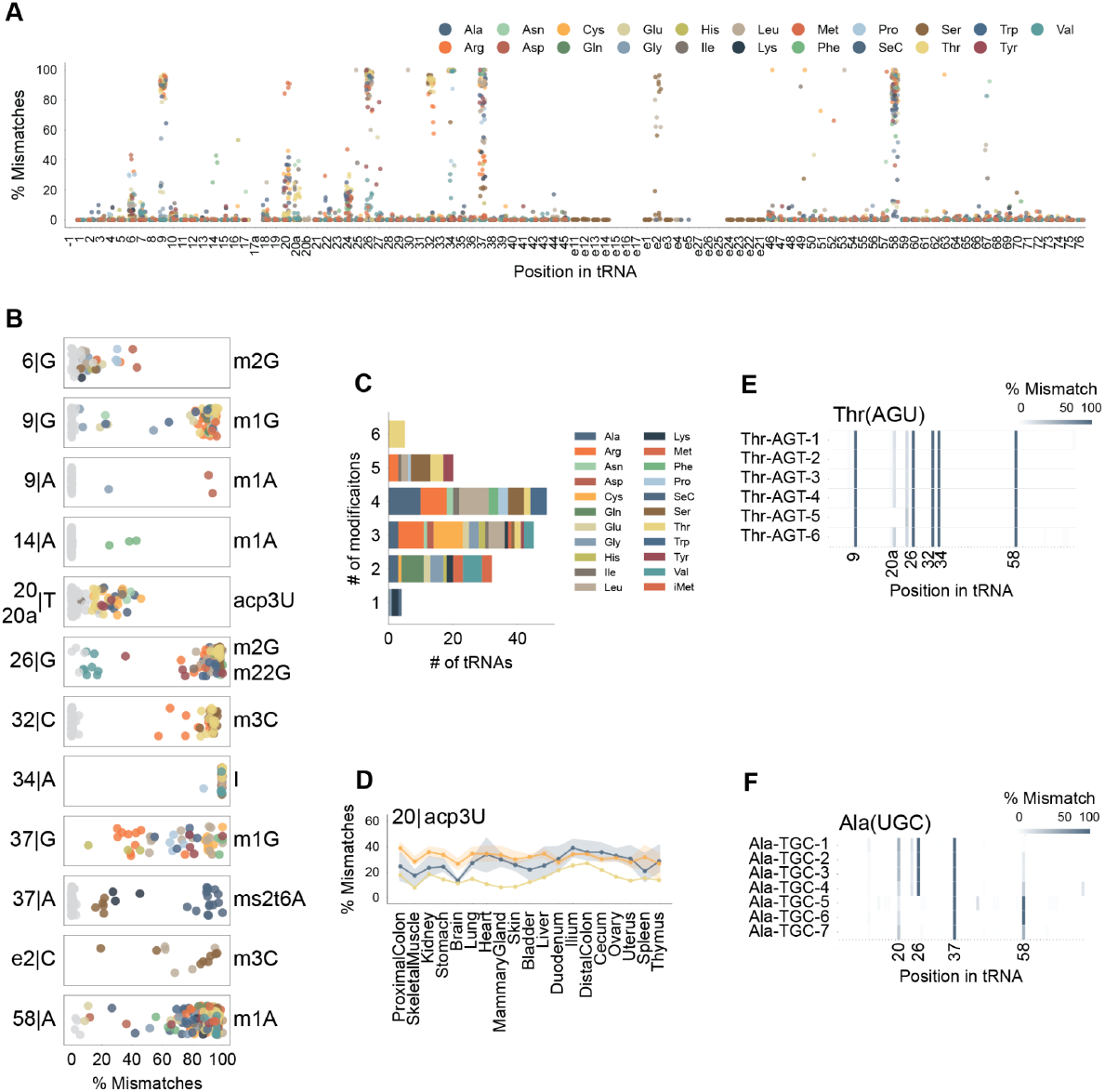
Nucleotide-resolution mapping of tRNA modifications reveals isodecoder-specific epitranscriptomic variation. **(A)** Landscape of modification-induced misincorporations. The strip plot shows the median percentage of mismatched bases at each position across all tRNA isodecoders with sufficient read depth (≥10 reads) in all tissues. Colors represent the tRNA isotypes. Known sites of tRNA modifications are highlighted where mismatches were observed. **(B)** Detailed view of misincorporation rates for specific known tRNA modification sites. Each strip plot displays the percentage of mismatches for a particular modification at a specific position, illustrating the variability across different tRNA isodecoders. **(C)** Number of modifications detected in each of the 155 detected tRNA isodecoders. **(D)** Tissue-specific variability of acp^3^U modification at position 20. The strip plot shows the percentage of mismatches for acp^3^U19 across different tissues. **(E-F)** Isodecoder-specific modification profiles. Bar plots display the percentage of mismatches at key modification sites for different isodecoders within the same anticodon family, revealing distinct epitranscriptomic patterns. **(E)** Modification profiles for Thr^(AGU)^ isodecoders. **(F)** Modification profiles for Ala^(UGC)^ isodecoders, highlighting differences, such as the loss of m^22^ G26 in tRNA-Ala-TGC-6.

We next investigated the differences of modification levels across the mouse tissues by comparing the RT-based misincorporation rates. Surprisingly, the vast majority of the tRNA modifications are highly consistent across all tissues, with only subtle differences observed. acp^3^U was one of the few modifications that fluctuated, with higher levels in the colon-related tissues, and slightly lower rates in brain and skeletal muscle (Figure 3D).

On the other hand, we observed notably different modification profiles of tRNA isodecoders with the same anticodon. Some of these tRNA isodecoders were found to be tissue-biased (Figure 2C). For instance, five out of six expressed tRNA^Thr(AGU)^ isodecoders carried acp^3^U at position 20a, whereas the heart-biased tRNA-Thr-AGT-5 has a C20a which prohibits it to have the same modification (Figure 3E). Interestingly, acp^3^U at this position has been shown to serve as an attachment site for N-glycans and may alter the glycoRNA profile of the heart tissue (Flynn et al. 2021). The lack of acp^3^U suggests that tRNA-Thr-AGT-5 may be involved in regulating glycoRNA in the heart. As mentioned previously, tRNA-Ala-TGC-5, 6, 7 were biased in the brain, with tRNA-Ala-TGC-6 being most abundant instead of tRNA-Ala-TGC-2/3 as in the other tissues. These three isodecoders have an A at position 26, resulting in a loss of m^2^_2_ G that was found in the other isodecoders (Figure 3F). More surprisingly, a compensatory increase in the m^1^A58 modification was also observed in these three isodecoders, indicating a diverse regulatory mechanism. These data highlight that not only do sequence variations exist amongst isodecoders, but also the epitranscriptomic modifications they harbor. Although these tRNAs have the same decoding feature, the differences in the base modifications may alter their thermostability and/or interactions with post-transcriptional machinery (Oerum et al. 2017). These variations indicate vastly diverse regulatory mechanisms that call for further investigation.

### Differential tDR pools elucidate potential tissue-relevant regulatory factors

With the identification of the tissue-biased mature tRNAs (Figure 2B), we hypothesized that these transcripts might serve as key precursors of tRNA-derived small RNAs (tDRs). To investigate this, reads < 65nt in length were isolated and categorized based on their mapping to each tRNA isodecoder: 5’ tDRs (derived within 10nt of the tRNA 5’ end), 3’ tDRs (derived within 10nt of the tRNA 3’ end), or internal tDRs (derived from regions not within 10nt of either end). Surprisingly, the tDR categories varied considerably across tissues (Figure 4A). For example, duodenum, spleen, and uterus predominantly featured 5’ tDRs while many other tissues including brain, heart, and ovary had a significant proportion of 3’tDRs. In contrast, the cecum and thymus displayed a more balanced distribution of 5’ and 3’ tDRs (Figure 4A). Some tissue-specific tDRs were derived from tissue-specific ‘minor’ isodecoders, including both a 5’ and 3’ tDR from brain-biased tRNA-Ala-TGC-6 (Figure 4B) and a 3’ tDR derived from the skin-biased tRNA-Ser-GCT-5 (Figure 4C), both of which contain key nucleotide substitutions compared to their corresponding ‘major’ isodecoder (Figure 2G). Another more regulated example was the tDRs derived from tRNA-Val-AAC-4 (Figure 4D). They were highly abundant in the duodenum; however, nearly absent in all other tissues despite tRNA-Val-AAC-4 being relatively more abundant in the brain amongst all tissues (Figure 2C). This suggests that not only the abundance of an isodecoder facilitates processing into tDRs, but other mechanisms, such as base modifications and/or variations in locally available enzymatic machinery, may play key roles in tDR synthesis.

**Figure 4.**
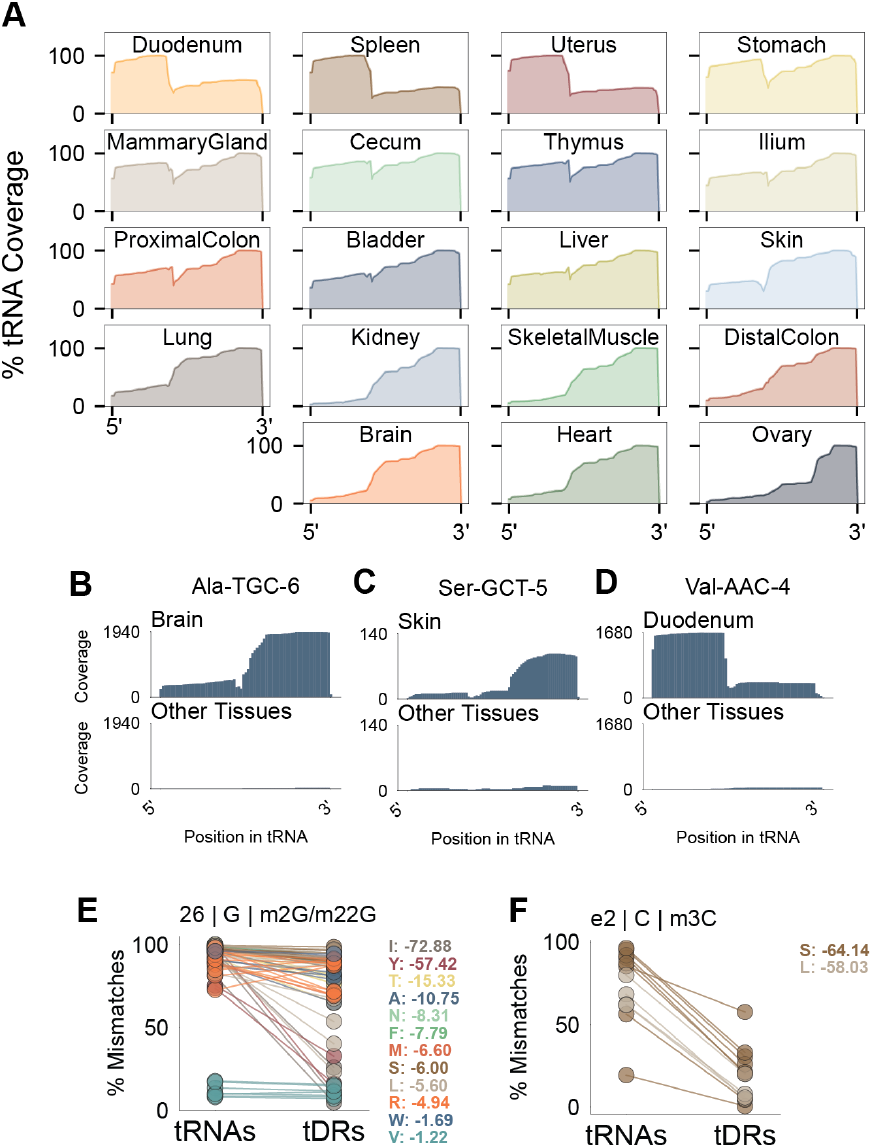
Tissue-specific processing tRNA-derived fragments (tDRs). **(A)** Meta-coverage profiles of tRNA-mapping reads for representative tissues. The plots illustrate the percentage of reads covering each position of a consensus tRNA, revealing tissue-specific cleavage patterns, such as the predominance of 5’ tDRs in the duodenum and spleen versus full-length tRNAs in the brain. **(B-D)** Coverage plots showing examples of highly tissue-specific tDRs. The plots compare the read coverage for a specific tRNA-derived fragment in its most abundant tissue versus the average of all other tissues. **(B)** Brain-specific tDRs derived from tRNA-Ala-TGC-6. **(C)** Skin-specific tDRs derived from tRNA-Ser-GCT-5. **(D)** Duodenum-specific tDRs derived from tRNA-Val-AAC-4. **(E-F)** Comparison of modification rates between parent tRNAs and their corresponding tDRs. The plots show the percentage of modification-induced mismatches, indicating that tDRs are generally hypomodified. **(E)** Mismatch rates for m^2^G/m^2^_2_G modifications at position 26. **(F)** Mismatch rates for the m^3^C modification in the variable loop (position e2).

To further understand the regulation of tDRs, we compared their modification rates with the mature tRNAs. Consistent with previous work, we found that tDRs had significantly fewer modifications compared to their precursors (Blaze et al. 2021) (Figures 4E and 4F). However, the number of modification losses varied across tDRs and the modification sites. For instance, we observed a dramatic ∼60-70% reduction of m^2^G/m^2^_2_G at position 26 in 5’ tDRs from Ile and Tyr tRNAs, but only a modest ∼5-10% reduction in those from Ala tRNAs (Figure 4E). In contrast, the variable loop modification m^3^C showed a more consistent, substantial reduction across Ser and Leu tRNAs (Figure 4F).

Collectively, these findings posit the generation of tDRs as an actively regulated, tissue-specific process, wherein the unique sequence and modification landscapes of tRNA isodecoders likely contribute to their cleavage susceptibility. The distinct tDR signatures discovered here substantially broaden the repertoire of potential regulatory small RNAs, strongly implicating them in shaping tissue identity, regulating physiological functions, and orchestrating responses to environmental stressors. Future investigations to identify the precise enzymatic machinery responsible for these cleavage events and to elucidate the specific biological roles of these tissue-biased tDRs will further the understanding of tRNA biology.

## Discussion

In this study, we have generated the most comprehensive collection and comparison of tRNA and tDR expression, along with their inferred RNA modifications, across 19 mouse tissues. These data expand on the finding that the majority of expressed tRNA isodecoders exhibit tissue-biased abundance, with many showing nearly exclusive tissue specificity. The sequence variations in tissue-biased tRNA isodecoders may lead to the loss or change of RNA modifications as illustrated in our findings that can alter the thermodynamic stability, folding kinetics, and affinity for interacting proteins, including aminoacyl-tRNA synthetases and modification enzymes (Westhof et al. 2022). These results may suggest a specialized “translational program” in individual tissue types that involves the regulation of the tRNA transcript abundance to optimize the expression of tissue-specific proteins or to serve as reservoirs for tDR synthesis, implying a layer of translational tuning at the level of individual tRNA isodecoders.

This interplay between the isodecoder sequence, modification status, and fragmentation is central to the biogenesis of tDRs. Our data show that tDR profiles are highly tissue-specific, often coinciding with the enrichment of their precursor tRNAs, as seen with tDRs derived from the brain-biased tRNA-Ala-TGC-6. However, precursor abundance is not the sole determinant, as shown by the duodenum-specific tDRs from tRNA-Val-AAC-4, a tRNA that is not specifically enriched in that tissue. This points to a more complex regulatory network where the presence of locally available enzymes and the epitranscriptomic state of the tRNA are more influential. The fact that tDRs have significantly fewer modifications compared to their full-length counterparts supports a model where specific modifications act as protective marks that inhibit cleavage as shown in previous studies.

While our study provides a more complete view of the tRNA pool across mammalian tissues, some limitations still exist. The modification mapping, based on reverse transcriptase misincorporation, is robust for select modifications. However, it cannot detect all modification types, particularly those that do not block or alter reverse transcription, such as m^5^C and pseudouridine, without additional chemical treatments, mass spectrometry, or nanopore direct RNA sequencing method. Furthermore, although our findings establish strong correlations between tRNA profiles and tissue identity, functional experiments such as isodecoder-specific knockouts or targeted modulation of tRNA-modifying enzymes will be required to better establish their biological roles.

In conclusion, we demonstrate that the tRNA pool is not a static component of the translational machinery but a dynamically regulated system that contributes to tissue identity and is profoundly altered in disease. This study provides a wealth of novel, tissue-specific biomarker candidates and uncovers numerous instances of isodecoder-specific regulation that open new avenues for investigating tRNA biology in health and disease.

## Methods

### RNA isolation and PNK treatment from mouse tissues

Tissues were harvested from three 6-8 week old wildtype female C57BL/6J mice at 2-3pm to best account for 12 hr light-dark cycles (7am-7pm light and 7pm-7am dark), which impact gene and protein expression during circadian variations. Total RNA was isolated from ∼15-100mg of whole bladder, brain, cecum, distal colon, duodenum, heart, ileum, kidney, liver, lung, mammary gland, ovary, proximal colon, skeletal muscle, skin, spleen, stomach, thymus, and uterus using 1mL of TRI Reagent and a tissue homogenizer, typically yielding ∼50-100ug of total RNA. Due to low RNA recovery for spleen and thymus, libraries were generated for only two of the three mice for these tissues. RNA 3’ de-phosphorylation was carried out as previously described (Huppertz et al., 2014) using 500ng of total RNA from each sample. Briefly, samples were treated with T4 Polynucleotide Kinase (T4PNK; New England Biolabs) in a modified 5x reaction buffer (350 mM Tris-HCl, pH 6.5, 50 mM MgCl2, 5mM dithiothreitol) under low pH conditions in the absence of ATP for 30 mins.

### OTTR-seq library preparation

OTTR-seq libraries were generated as previously described (Upton et al. 2021). Briefly, total PNK-treated RNA was 3’ tailed using mutant BoMoC RT in buffer containing only ddATP for 90 minutes at 30°C, with the addition of ddGTP for another 30 minutes at 30°C. This was then heat-inactivated at 65°C for 5 minutes, and unincorporated ddATP/ddGTP were hydrolyzed by incubation in 5 mM MgCl2 and 0.5 units of shrimp alkaline phosphatase (rSAP) at 37°C for 15 minutes. 5 mM EGTA was added and incubated at 65°C for 5 minutes to stop this reaction. Reverse transcription was then performed at 37°C for 30 minutes, followed by heat inactivation at 70°C for 5 minutes. The remaining RNA and RNA/DNA hybrids were then degraded using 1 unit of RNase A at 50°C for 10 minutes. cDNA was then cleaned up using a MinElute Reaction CleanUp Kit (Qiagen). To reduce adaptor dimers, cDNA was run on a 9% UREA page gel, and the size range of interest was cut out and eluted into gel extraction buffer (300mM NaCl, 10mM Tris; pH 8.0, 1mM EDTA, 0.25% SDS) and concentrated using EtOH precipitation. Size-selected cDNA of ∼15-100 nt was then PCR amplified for 12 cycles using Q5 High-fidelity polymerase (NEB #M0491S). Amplified libraries were then run on a 6% TBE gel, and the bands corresponding to inserts of ∼15-100 nt the size range of interest were extracted to further reduce adaptor dimer inclusion. Gel slices were eluted into gel extraction buffer (300mM NaCl, 10mM Tris; pH 8.0, 1mM EDTA) followed by concentration using EtOH precipitation. Final libraries were pooled and sequenced on an Illumina NextSeq 500 with 150-cycle high-output kit.

### Sequencing Data Processing

Sequencing adaptors were trimmed from raw reads using cutadapt v1.18 (Martin 2011) [--no-indels -O 15 -m 15 -a GATCGGAAGAGCACACGTCTGAACTCCAGTCAC], then UMI’s were extracted using umi_tools (Smith et al. 2017) [extract --extract-method=string --bc-pattern=NNNNNNN], finally followed by the removal of a single base added during library preparation from the 5’ end using cutadapt [-u -1 -q 10]. To address the variable incorporation of mature tRNAs and tDRs across tissues, these reads were separated into “mature tRNAs” or “tDRs” based on read lengths of >=65 and <65nt, respectively. Reads were subsampled from each tissue sample to account for differences in sequencing depth to 200,000 reads per sample for those >=65nt and 1,000,000 reads per sample for those <65nt using seqtk [sample -s100]. These were then sequentially aligned using bowtie2 (Langmead and Salzberg 2012) [--very-sensitive -k 100] to vector contaminants (UniVec), rRNAs, high confidence tRNAs (with the addition of the -CCA tail) of mouse genome GRCm38/mm10 obtained from GtRNAdb (Chan and Lowe 2016), miRNAs from miRbase (Kozomara et al. 2019), snRNAs/snoRNAs/lncRNAs from Ensembl (Dyer et al. 2025), processed mRNA transcripts (protein_coding) from Ensembl (Dyer et al. 2025), and lastly to the GRCm38/mm10 mouse reference genome. Counts associated with these features were then quantified choosing the feature annotation with the largest degree of overlap. Multimapping ncRNA reads with more than one highest-scoring alignment were randomly selected. The resulting count data underwent normalization and differential expression analysis using DESeq2 (Love et al. 2014). Visualizations were generated using custom Python scripts.

### Tau tissue-specificity score

The tau value for all transcripts were calculated using the following equation as previously described (Kryuchkova-Mostacci and Robinson-Rechavi 2017):

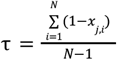

where *N* is the total number of tissues assessed, and x_j,i_ is the expression value of tissue *i* normalized by the maximal expression of any tissue for transcript *j*. Based on previous work (Kryuchkova-Mostacci and Robinson-Rechavi 2017), a value of tau >= 0.85, is expressed in primarily one tissue, values of 0.35-0.85 are indicative of transcripts biased to groups of tissues (intermediate), and values less than 0.35 as considered equally abundant across all tissue types.

### Modification-induced misincorporations

To determine the potential tRNA modification sites present in the tRNAs and tDRs, a custom python script was used. Briefly, tRNA/tDR transcripts were summed across tissues and filtered to have at least 10 non-normalized reads along with >50% of these being uniquely mapped to an isodecoder. The counts of adenines, guanines, cytosines, thymines, and deletions mapped to the individual nucleotides of each isodecoder were then summed, and the number of non-reference bases was calculated. By dividing this value by the total number of counts for that nucleotide, multiplied by 100, a percent mismatch was determined. A confidence score was calculated using a previously described Bayesian model (Bahn et al., 2012; Li et al., 2008) based on the number of supporting reads and the percent edited reads to determine whether or not a base modification was present. Modification sites with less than 5% of mismatched reads and a 95% confidence score were not considered modified. Downstream data visualizations were generated with a custom python script using these mismatch values for each tRNA or tDR across each tissue.

## Data Access

The sequencing data for this study can be accessed at GSE309058.

## Acknowledgments

We thank Lucas Ferguson and Kathleen Collins at University of California, Berkeley for providing reagents for OTTR-seq experiments, and their considerable assistance and consultation in optimizing the protocol for tRNAs. This work was supported by a grant from the National Human Genome Research Institute, National Institutes of Health [R01HG006753 to T.L.].

## Authors’ Contributions

AM and TL conceptualized the project. AM performed the sequencing experiments and analyzed the data. All authors wrote and edited the manuscript. The authors read and approved the final manuscript.

